# Sugar units, not chilling units, control endodormancy duration in plants – potato tuber as a case study

**DOI:** 10.1101/2021.05.10.443356

**Authors:** Raz Danieli, Shmuel Assouline, Bolaji Babajide Salam, Paula Teper-Bamnolker, Eduard Belausov, Yael Friedmann, David Granot, Dani Eshel

## Abstract

Endodormancy (ED) is a crucial stage in the life cycle of many perennial plants, regulated by genetic and environmental factors. Chilling units, growth regulators, and nutrient supply are considered inducers of ED release, but the mechanism governing ED duration is poorly understood. The potato tuber has been used as a model system to study metabolic processes associated with ED release. Cold-induced sweetening is a well-known response of the tuber to chilling. Here, we found that cold stress induces an increase in sugar units in association with plasmodesmatal closure in the dormant bud cells. Tuber sweetening was associated with shortened ED duration after cooling. Heat exposure also caused sugar unit accumulation followed by faster ED release. A logistic function was developed to predict ED duration based on sugar unit measurements. We discovered that ED release is better correlated with the accumulation of sugar units compared to chilling units. CRISPR/Cas9 knockout of the vacuolar invertase gene (*StVInv*) induced longer ED, but only in cultivars in which the mutation modified the level sugar units. Our results suggest that sugar units are better predictors of vegetative bud ED duration than chilling units.

## INTRODUCTION

The ability to form buds and to undergo cycles of growth and dormancy has been central to the evolution of the perennial life strategy. Perennial plants have developed a dormancy mechanism in temperate regions to overcome severe cold temperature conditions and frost. Lang et al.^1^ distinguished three types of bud dormancy: endodormancy (ED), maintained by internal bud signals; paradormancy, promoted by other plant organs; and ecodormancy, induced by environmental factors. It is unclear whether the genetic mechanisms controlling these different dormancy types are related, or whether diverse endogenous and ecological stimuli trigger signaling pathways that converge to activate a joint genetic program. Most deciduous tree species become dormant in response to decreasing temperature. ED release depends on the chilling requirement, i.e., the number of chilling units to which the trees are exposed during the winter ^2^. The number of chilling units required for bud break during the spring varies among cultivars and species ^3–5^. Accordingly, chilling units are commonly used to predict dormancy duration ^6,7^.

Potato tubers have been used as a model system to study ED characteristics ^8–11^. Tuber dormancy represents a case in which an otherwise annual plant generates a vegetative perennating organ with an unclear chilling requirement ^3,12^. The potato tuber is a stem formed by swelling of the subapical underground stolons ^13^. As the tuber elongates, a growing number of lateral bud meristems (termed eyes) are formed in a spiral arrangement on its surface ^14^. After harvest, tuber buds are generally dormant and will not sprout or grow, even if the tubers are placed under optimal conditions for sprouting ^15,16^. This dormancy is defined as ED ^1^, and is due to an unknown endogenous signal(s) that mediates suppression of tuber-bud meristem growth ^17^. Tuber dormancy is thought to be a physiological adaptation to intermittent periods of environmental limitations. Therefore, it is a survival mechanism that prevents sprouting when the potato plant would be exposed to extreme environmental conditions ^18^. The duration of the tuber ED period is primarily dependent on the genotype, crop growth, and postharvest conditions ^19–22^. Following a transition period of between 1 and 15 weeks, depending on the storage conditions and variety, dormancy is released, and the tuber apical bud starts to grow ^20^.

Buds are actively growing plant organs that serve as strong sinks for sugars to meet their metabolic demands and support their growth. The bud’s growth capacity depends on its sink strength in terms of its ability to acquire and use available sugars. Thus, buds have to compete for sugars, which constitute the main source of carbon and energy ^11,23–25^. The association between bud outgrowth and mobilization of starch reserves in stem tissues has been well-documented, especially in perennial plants ^26,27^. High activity of sugar-metabolizing enzymes leads to increased sugar absorption by the bud ^24,27–30^.

During tuber development, the storage parenchyma converts soluble assimilates (sucrose, amino acids) into polymeric reserves (starch and storage proteins) ^31,32^. At maturity, over 70% of the tuber carbohydrate reserves are sequestered as starch, which must be converted into transport-compatible solutes for sprouting initiation and growth ^33,34^. Sucrose synthesis appears to be a dominant anabolic pathway in the dormant tubers’ parenchyma ^35^. The phenomenon of cold-induced sweetening, resulting from the accumulation of reducing sugars in cold-stored potato tubers, has been mainly studied for its impact on potato processing ^36-38^. During cold-induced sweetening, sucrose synthesis increases and some sucrose is transported to the vacuole, where it is hydrolyzed to glucose and fructose ^37,39^. This step is predominantly controlled by vacuolar acid invertase (VInv), an enzyme that is strongly associated with the accumulation of reducing sugars during cold storage ^40,41^. Silencing of the potato *StVInv* gene results in effective control of cold-induced sweetening ^42,43^.

Although the requirement of a chilling period for ED release is well-established in vegetative buds, such as tuber buds, the mechanism by which cold stress shortens ED duration is not clear. Phytohormones regulate bud dormancy in potato tuber ^17^. Gibberellins, cytokinins and brassinosteroids induce dormancy release and promote tuber sprouting ^44,45^, while ethylene and abscisic acid (ABA) promote the maintenance of dormancy ^46^. To encourage growth, soluble sugars, and other growth-promoting signals, need to move through the symplastic pathway into the dormant bud ^47^. During the dormancy period, symplastic intercellular communication through plasmodesmata (PD) is blocked by callosic dormancy sphincters; it is reestablished upon dormancy release ^47,48^. Viola et al. (2007) showed differential sugar metabolism in dormant and growing buds of potato tubers, suggesting blockage of their transport to the dormant bud.

In this study, we used dormant potato tuber as a model system to determine the role of sugar accumulation, induced by temperature stress, in determining ED duration of vegetative buds. We establish a role for total sugar units as a better predictor of ED duration than chilling units.

## RESULTS

### Cold-induced sweetening activates early ED release

To study how cold stress regulates potato tuber ED, freshly harvested ‘Desiree’ tubers were stored at 14°C for 2 weeks for curing and conditioning. They were then transferred to 4, 6, 9, or 12 °C, or left at 14°C, for 4 weeks. Tubers were returned to 14°C to induce sprouting, a sign of ED release. Every 2 weeks, five tubers from each treatment were sampled for sugar analysis. As expected, cold stress induced elevated levels of sucrose, glucose and fructose (Figures 1A–1C). Sucrose concentration peaked at 2 weeks of cold storage, followed by peaks of glucose and fructose, which accumulated mainly in tubers that were stored at temperatures below 6°C (Figures 1B and 1C).

**Figure 1.**
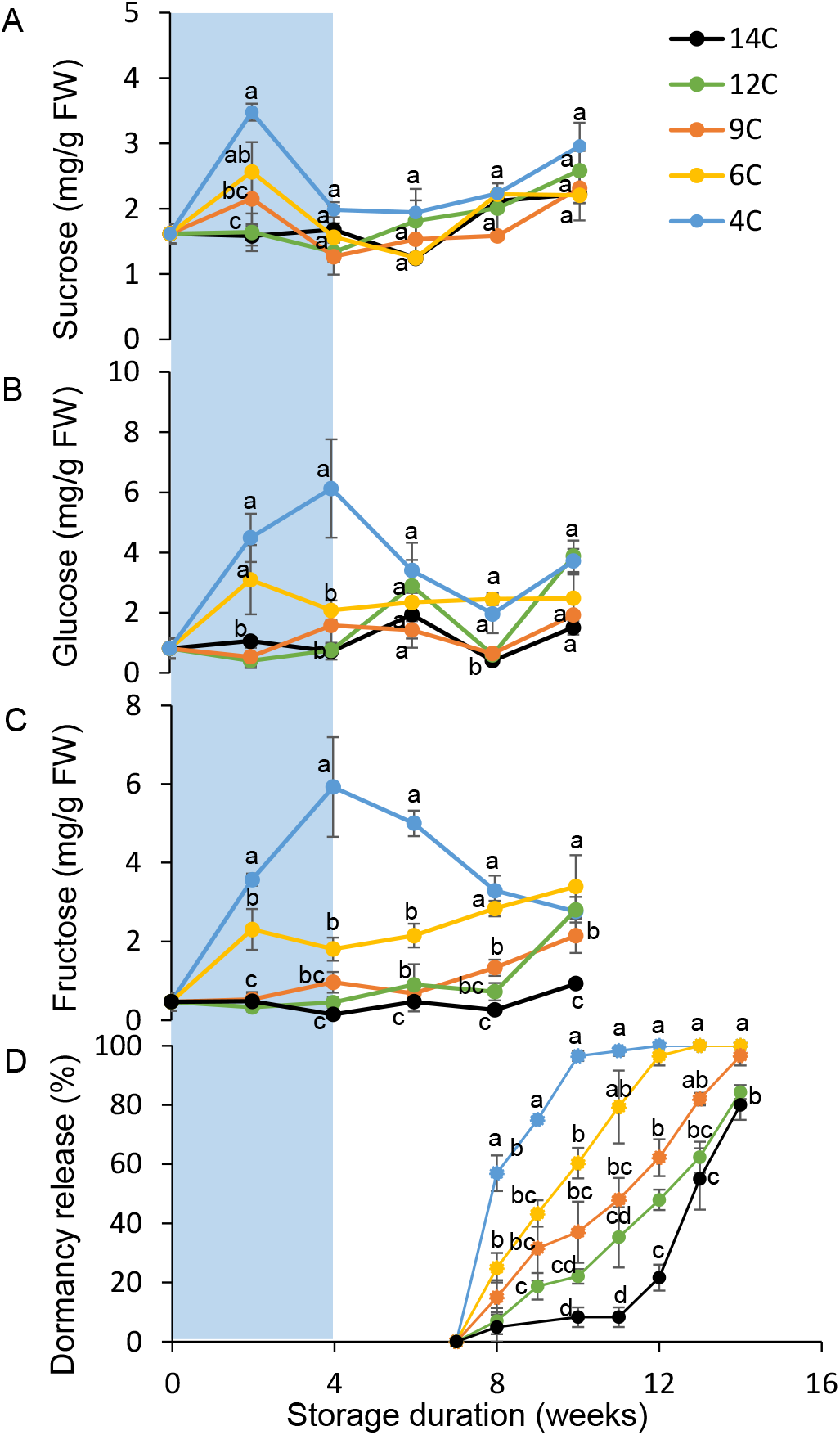
Exposure to cold storage temperature induces early endodormancy release. ‘Desiree’ tubers were stored for 4 weeks (light blue background) at five different temperatures (14, 12, 9, 6, and 4°C) and then transferred to 14°C storage. Every 2 weeks, five tubers from each treatment were sampled for (A) sucrose, (B) glucose, and (C) fructose quantification. (D) In each treatment, three repeats of 20 tubers were measured every week for dormancy release. A tuber was considered dormant until at least one bud reached 2 mm in length. Error bars represent SE. Different letters indicate significant differences between treatments at each time point (one-way ANOVA, *p* < 0.05).

Surprisingly, 4°C storage for 4 weeks was most efficient at shortening ED duration, with just 9 weeks until 75% of the tubers sprouted (Figure 1D). Exposure to higher temperatures induced longer dormancy, up to 14 weeks for storage at 14°C (Figure 1D). A similar pattern was found after 8 weeks of cold exposure (Figure S1); however, sprouting (dormancy release) differences between treatments were reduced, suggesting suppression of growth by the cold temperature after ED release, namely, ecodormancy.

The ED duration data (Figure 1D) presented a sigmoidal behavior that was well-represented by the logistic function:

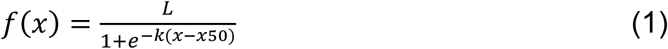

where *L* represents 100% dormancy release, *x*_50_ is weeks until dormancy of 50% of the tubers is released, and *k* is the dormancy-release rate. We fitted the function to the measured ED duration data for the different conditions and characterized the *x*_50_ and *k* parameters that led to the best fit for each case. The agreement between the observed percentage of dormancy release and that predicted using the logistic function was very high (Figure S2A). The parameter *x*_50_ decreased dramatically in the first 4 weeks and then increased gradually with prolonged cold exposure (Figure S2B). The *k* value, i.e., dormancy-release rate, showed a slight and gradual increase up to 8 weeks, followed by a steep increase for longer cold exposure (Figure S2C). We suggest that ED duration following exposure to 4°C for 4 weeks is limited to the first 8 weeks of storage, as suggested by the *k* value.

Tuber exposure to 2 weeks, instead of 4 weeks, at 4°C, followed by their transfer to 14°C, showed lower sugar accumulation and longer ED duration, suggesting the need to absorb a sufficient amount of chilling units for ED release (Figure 2). No bud burst was observed in tubers that were continuously exposed to 4°C (16 WC treatment in Figure 2), suggesting that ED release occurs at cold temperature, but higher temperature is needed for bud growth. We obtained similar results in two other potato cultivars—Sifra, and Tyson (Figure S3), suggesting a more general phenomenon of shortened ED period due to cold stress followed by parenchyma sweetening.

**Figure 2.**
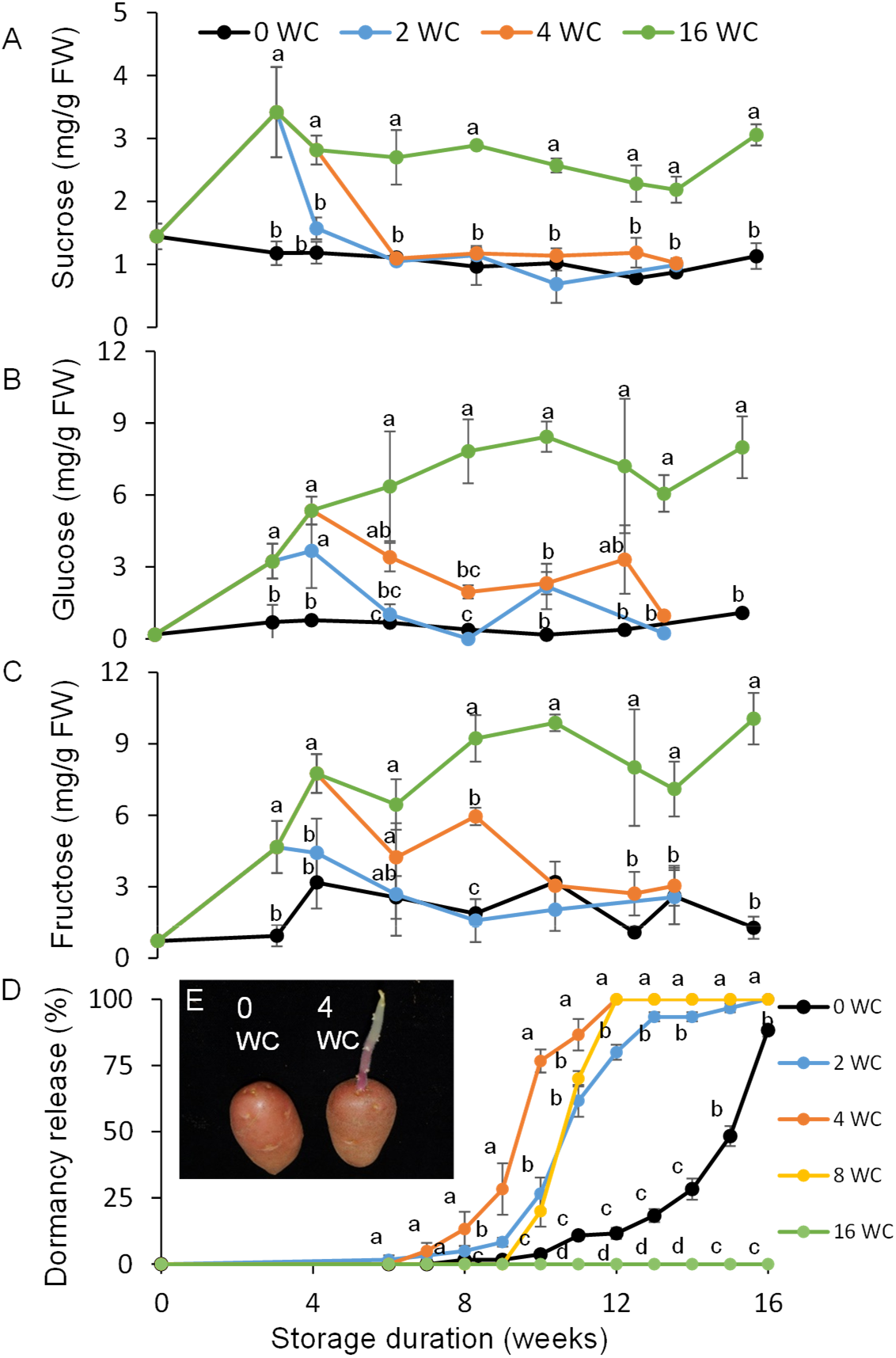
A sufficient amount of chilling units is needed for endodormancy release. ‘Desiree’ tubers were exposed to five cold (4°C) treatments of different duration (0, 2, 4, 8 and 16 weeks; WC) and then transferred to 14°C storage. Every 2 weeks, five tubers from each treatment were sampled for (A) sucrose, (B) glucose, and (C) fructose quantification. (D) In each treatment, three repeats of 20 tubers were measured every week for dormancy release. A tuber was considered dormant until at least one bud reached 2 mm in length. (E) Representative tubers after 15 weeks of storage. Error bars represent SE. Different letters indicate significant differences between treatments at each time point (one-way ANOVA, *p* < 0.05).

### Symplastic-connection blockage induced by cold stress delays ED release during parenchyma sweetening

A high level of soluble sugars, induced by 4 weeks of cold stress, did not induce bud burst immediately after transferring the tubers to optimal growing conditions. Dormancy was released 4 weeks later, when the soluble sugars had dramatically decreased. PD closure was determined as a possible cause for the time gap between sugar elevation and dormancy release. Tuber apical buds were sampled during and after cold stress and stained with Aniline blue. The buds were dissected and analyzed under a confocal microscope for callose accumulation in the cell-wall PD, which indicates blockage of the symplastic connection. We found an increase in the number of closed PD per cell during the first 4 weeks of cold stress (Figure 3). Terminating the cold exposure resulted in a gradual decrease in the number of closed PD per cell during the following 4 weeks of storage at 14°C (Figure 3), suggesting blockage of the cell-to-cell connection as a response to cold stress. We expect that such PD blockage reduces the transport of nutrients to the dormant bud, thereby delaying ED release.

**Figure 3.**
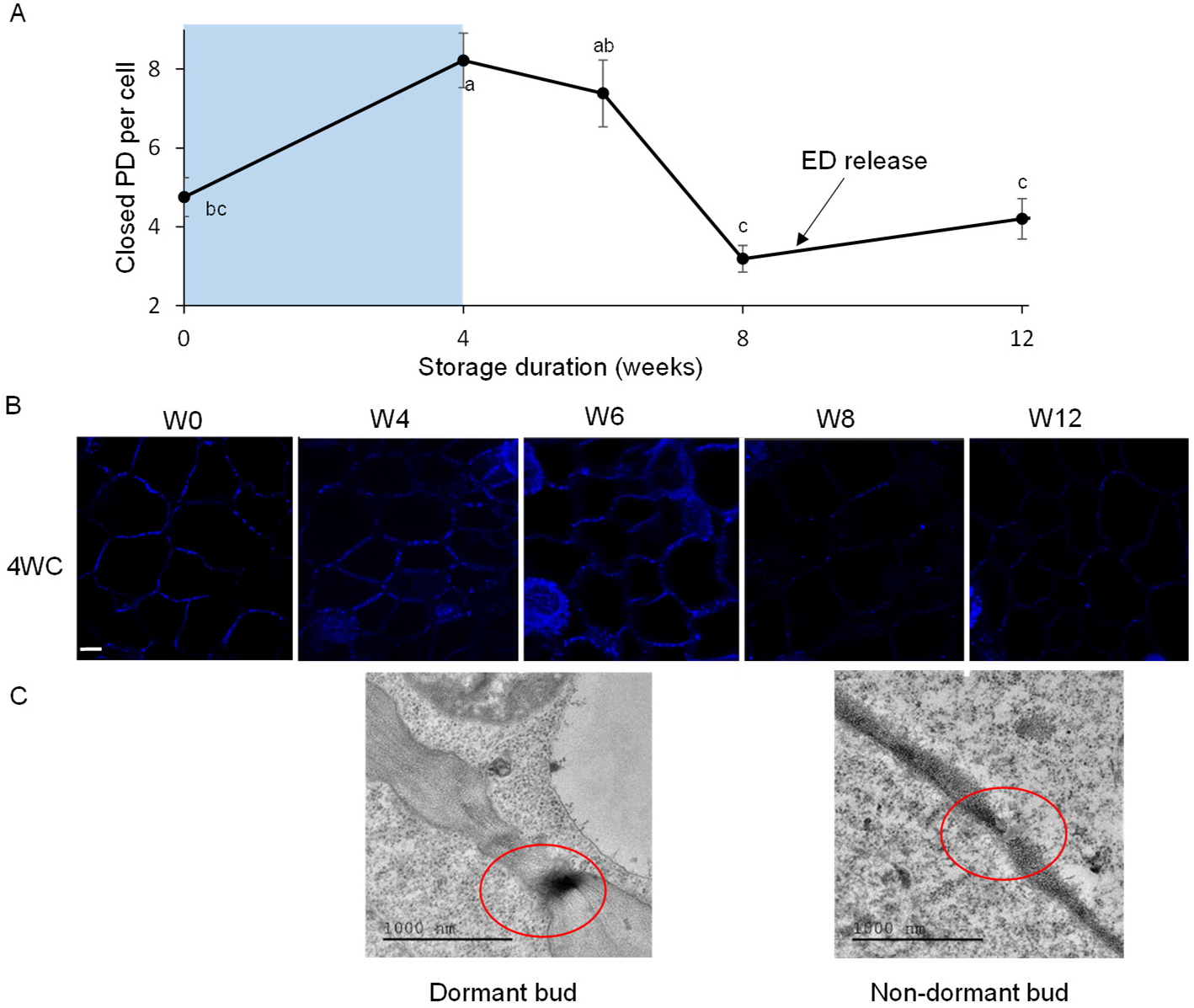
Symplastic connection is blocked during cold stress. ‘Desiree’ tubers were stored for 4 weeks at 4°C (light blue background; WC) and then transferred to 14°C storage. (A) Apical buds were sampled at 0, 4, 6, 8, and 12 weeks in storage and dipped in Aniline blue for callose staining. The amount of callose accumulation (indicating number of closed plasmodesmata [PD]) per cell was analyzed. Black arrow indicates *x*_50_ time point. Each time point represents 30 cells from four buds; different letters indicate significant differences between time points (one-way ANOVA, *p* < 0.05). (B) Representative confocal images at 0–12 weeks (W0–W12) in storage. Bar = 5 μm. (C) Representative transmission electron microscopy images of PD (circled in red) in dormant (left) and non-dormant apical bud. Bar = 1 μm.

### ED release is better associated with the accumulation of sugar units than chilling units

Chilling requirements vary among deciduous perennial tree species and are represented by the number of chilling units required for bud break during the spring ^3^. We tested whether this is the case in ED release of potato tuber buds. Defining 14°C as the ideal sprouting temperature, we counted each degree below 14°C on each day of cold exposure as 1 chilling unit. To our surprise, we found that chilling unit exposure correlated with *X*_50_ with an adjusted R-square value (R^2^ Adj) of only 0.56 (Figure 4A). Sucrose and glucose showed the highest correlation to *x*_50_, with R^2^ Adj of 0.76 and 0.79, respectively (Figures 4B and 4C). One sugar unit can be considered a lumped indicator of all sugar types, and can be estimated using the following equation:

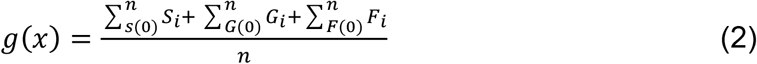

where *S, G* and *F* represent sucrose, glucose, and fructose, respectively, and *n* is the number of measurements during the indicated period (4 or 8 weeks). Sugar units gave a higher R^2^ Adj fit than chilling units (Figure 4D).

**Figure 4.**
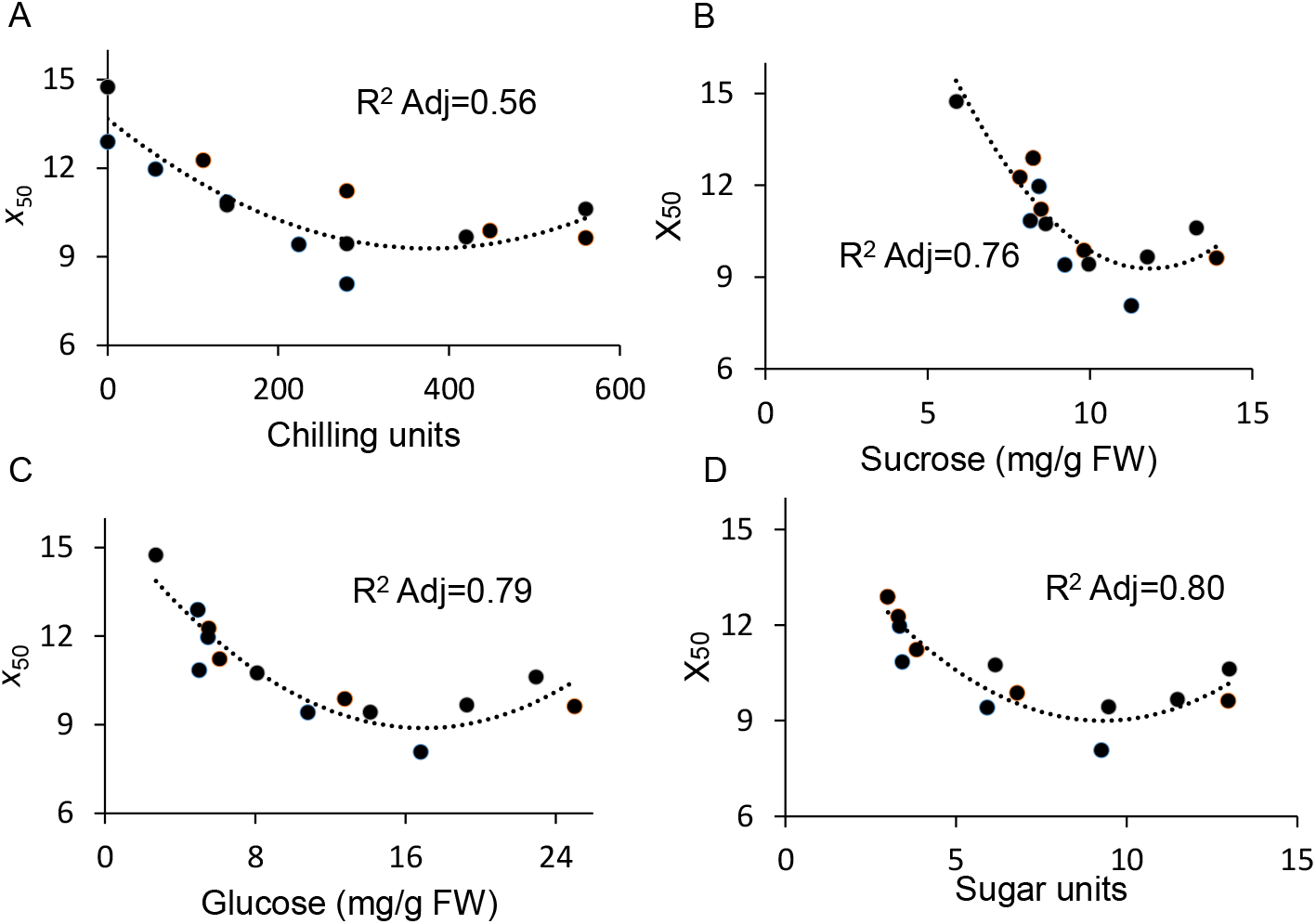
Dormancy release is more closely related to sugar units than chilling units. Polynomial order 2 correlation between the fitted logistic function parameter *x*_50_ and (A) chilling units, (B) sucrose and (C) glucose accumulation, and (D) sugar units from both the storage temperature experiment and cold duration experiment. R^2^ Adj was calculated with JMP.

To separate the sweetening process from the cold-stress effect, we induced early ED release by heat treatment, as done previously ^11^. ‘Desiree’ tubers were exposed to 0, 1, 2, 4, and 6 weeks of heat (WH) and then transferred to 14°C storage. Heat treatment induced sucrose accumulation in the tuber parenchyma and had a minor effect on hexose levels (Figures 5A–5C). Dormancy duration was shortest with 4 WH and longest with no heating (0 WH; Figures 5D and 5E). The parameter *x*_50_ was highly correlated to sucrose and sugar units and less correlated to glucose (Figures 5F–5H). The delayed-ED-release phenotype with 6 WH, together with the high value of *k*, suggested that ED release is followed by growth inhibition induced by heat (ecodormancy; Figures 5D and 5I).

**Figure 5.**
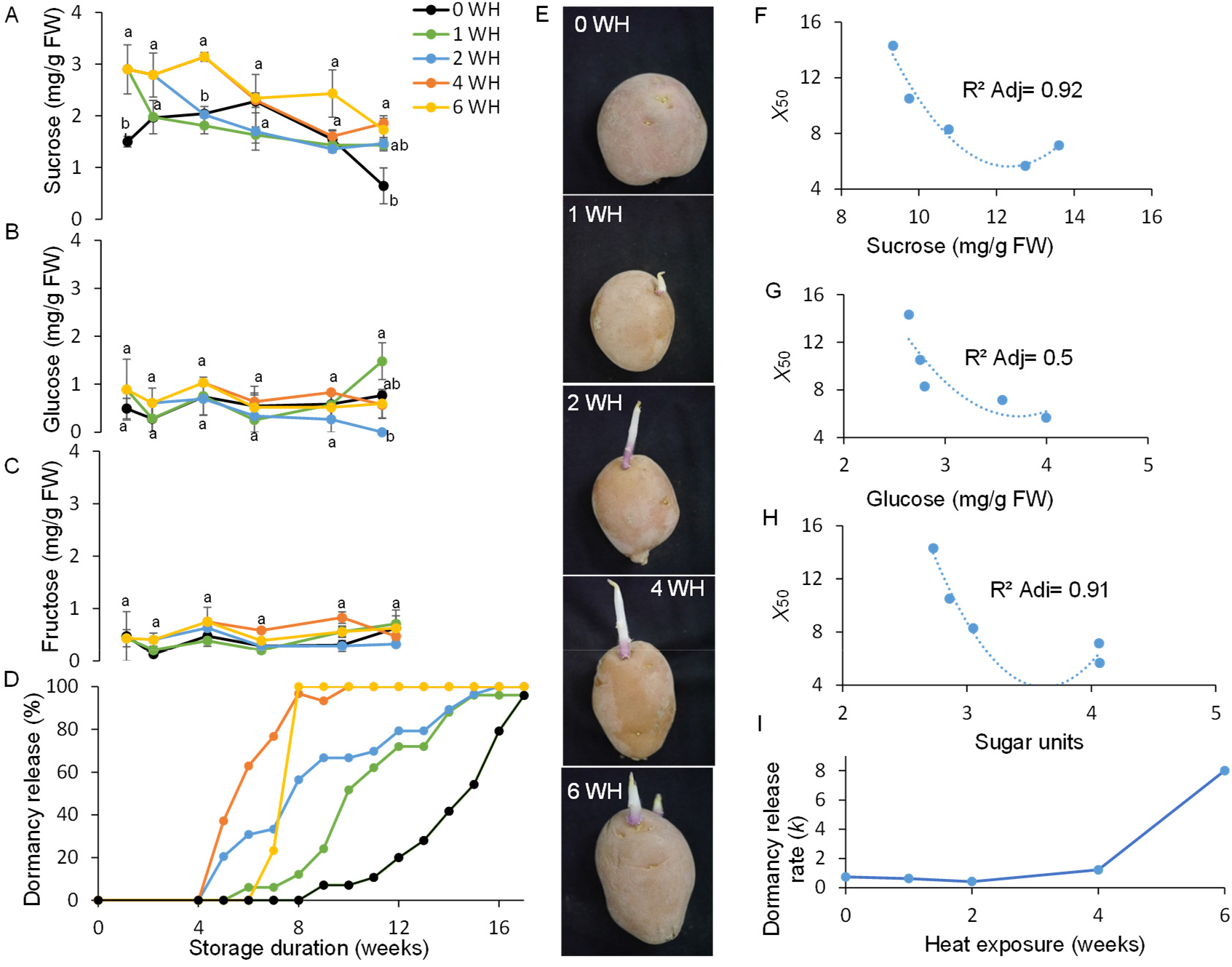
Endodormancy release induced by heat treatment is correlated with sugar unit accumulation. ‘Desiree’ tubers were exposed to 0, 1, 2, 4, and 6 weeks of heat (33°C; WH) and then transferred to 14°C storage. Every 2 weeks, four tubers from each treatment were sampled for (A) sucrose, (B) glucose and (C) fructose quantification. Error bars represent SE. Different letters indicate significant differences between treatments at each time point (one-way ANOVA, *p* < 0.05). (D) In each treatment, 30 tubers were measured every week for dormancy release. A tuber was considered dormant until at least one bud reached 2 mm in length. (E) Representative images of tubers after 13 weeks of heat treatment and storage. (F-H) Polynomial order 2 correlation between the fitted logistic function parameter *x*_50_ of sucrose (F), glucose (G) and sugar unit (H) level. (I) Values of the fitted logistic function parameter *k* for various heat-exposure durations.

### *StVInv-knockout* extends ED duration differentially and in association with modifications in sugar units

Sucrose and hexose accumulated in the tuber parenchyma of both ‘Desiree’ and ‘Brooke’ in response to cold exposure (Figure 6). ‘Desiree’ *StVInv-knockout* lines (#7 and #8) showed a reduction in sugar units following cold stress, associated with longer ED duration (Figure 6). In contrast, ‘Brooke’ *StVInv* lines (#29 and #60) showed no significant differences compared to the respective wild type in sugar unit accumulation or ED duration (Figure 6). These results suggested that total sugar units are the primary regulator of ED.

**Figure 6.**
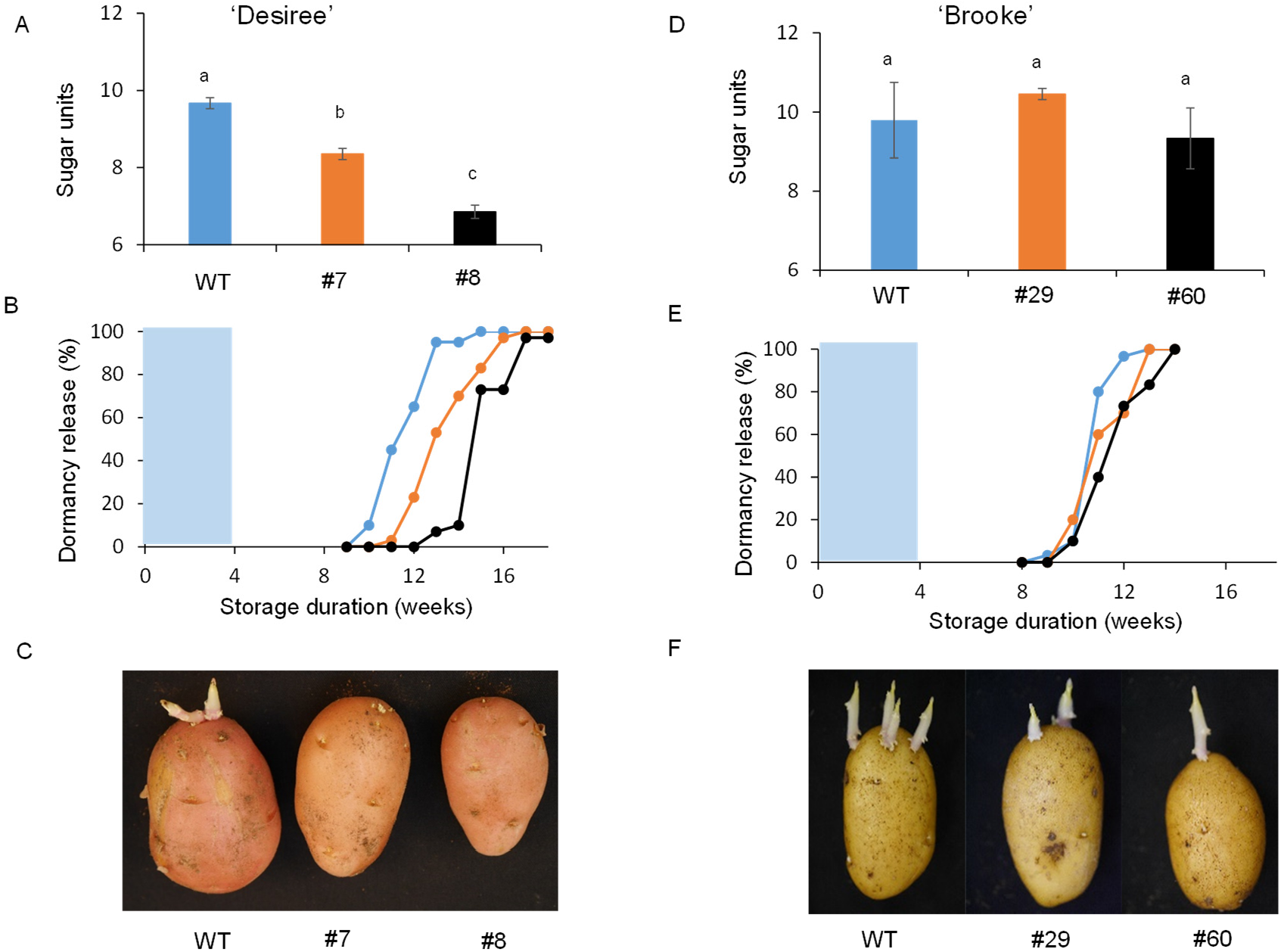
*StVInv-knockout* lines have modified endodormancy duration associated with the effect on sugar unit accumulation. Wild-type (WT) and *StVInv*-knockout tubers of cvs. Desiree (A–C) and Brooke (D–F) were stored for 4 weeks at 4°C (light blue background) and then transferred to 14°C. (A,D) Every 2 weeks, five tubers from each line were sampled for sugar (sucrose, glucose and fructose) quantification, and then sugar units were calculated. (B,E) Thirty tubers were tested every week for dormancy release. (C,F) Representative images of tubers after 13 weeks of cold stress and storage. Error bars represent SE. Different letters indicate significant differences between treatments (one-way ANOVA, *p* < 0.05).

## DISCUSSION

### Cold exposure induces early ED release of potato tuber

A cold-exposure requirement for early dormancy release has been shown in perennial tree buds ^49^ and bulb buds of *Gladiolus* ^50^, *Liliaceae* ^51,52^ and onions ^53^. Cold storage is widely used as a tool to delay the sprouting of postharvest potatoes, mainly by maintaining ecodormancy following ED release ^54^. On the other hand, studies have shown that temporary cold stress results in shorter dormancy in potato tubers ^55,56^. We found that both temperature level and chilling unit accumulation modify ED duration. In addition, we discovered that a logistic function accurately describes dormancy release in potato tubers (Figure S2). This equation emphasizes two important parameters: *x*_50_, which expresses the number of weeks until 50% of the tubers have undergone dormancy release and normalizes dormancy release data to one number, and *k,* which describes the dormancy-release rate. We found that the ED period is typically characterized by *k* < 2, whereas in ecodormancy, *k* > 8 (Figure S2). This enables distinguishing, mathematically, ED from the ecodormancy period. The logistic function has been suggested as a model for common knotgrass (*Polygonum aviculare*) seed dormancy release, and to predict bud break in sweet cherry (*Prunus avium*) trees ^57–59^. Here, we showed that chilling units induce early ED release of tuber buds (Figures 1 and 2) in a pattern similar to that in tree buds ^2^. These results suggest that the cold-requirement mechanism found in other plants also exists in potato tubers.

### ED release is better associated with the accumulation of sugar units than chilling units

Cold-induced sweetening is a well-studied process in which carbohydrates are converted from starch to soluble sugars, mainly sucrose, glucose, and fructose, due to cold stress ^37,60^. When ED is released and the temperature allows bud growth, sugars are consumed by higher respiration, starch resynthesis, or sprouting ^34^. Consequently, a specific threshold of soluble sugar availability probably needs to be crossed. Therefore, sugar units are calculated as the average soluble sugar availability during the ED period. Analysis of the dormancy duration data by means of the logistic function allowed identifying, quantitatively, the processes involved in the dormancy release, and partitioning between the generally used chilling units, and sugar units, introduced in this study. The parameter *x*_50_ showed a higher correlation with sugar units than with chilling units (Figure 4). The strong correlation between *x*_50_ and sugar units remained when ED release was induced by heat stress (Figure 5). We can assume that increasing sugar units in response to temperature stress is an osmoprotective or scavenging response ^61,62^.

VInv breaks sucrose into glucose and fructose; silencing *StVInv* in potato tubers results in a different pattern of cold-induced sweetening symptoms ^63,64^. Here, we exposed *stvinv* lines of two different cultivars to cold stress, and decreased sugar unit accumulation was only observed in ‘Desiree’ (Figure 6). Although both cultivars were exposed to the same cold stress, only ‘Desiree’ *stvinv* lines showed prolonged ED in association with the differential sugar units accumulation (Figure 6). RNAi silencing of *StVInv* in cv. Russet Burbank causes an increase in tuber sucrose, mostly in the parenchyma and not in the bud ^11^. We suggested that parenchyma sucrose induces signaling for multiple stems in the sprouting tuber, rather than energy mobilization from the parenchyma to the bud ^11^. Here, knockout of ‘Brooke’ *StVInv* did not affect sugar units and there was no change in ED duration. Silencing of potato alpha-amylase 23 (*StAmy23*) also inhibits dormancy release in cv. Solara ^65,66^. Those authors suggested that more extended dormancy is related to the shortage of reducing sugars. According to their results, *stamy23* lines had reduced glucose levels and no concordant difference in sucrose levels, suggesting lower sugar-unit formation during ED. Sugars have been suggested to regulate sprouting via the trehalose-6-phosphate (T6P)–sucrose pathway ^34^. Debast et al. ^67^ managed to influence tuber dormancy duration by altering T6P content in cv. Solara. They linked the dormancy phenotype with ABA content, and suggested that T6P regulation of SNF1-related protein kinase (SnRK1) might regulate ABA content. Here we suggest that sugar units are the best predictor of ED duration, probably as a result of serving as an energy source and signal for dormancy release.

### The time gap between sugar accumulation and ED release is associated with PD closure

Vegetative bud burst requires the formation of symplastic connections to the bud to allow metabolic flow prior to dormancy release ^35,68^. We found that sugar levels in the tuber parenchyma decrease dramatically when cold exposure is terminated. At the time of bud burst, the cold-stressed tubers’ sugar content does not differ from that of unexposed tubers (Figures 1 and 2). We showed that PD remain blocked during cold stress and 2 weeks after it, suggesting that the symplastic connection has yet to be formed (Figure 3). Viola et al. ^35^ demonstrated that sugars must reach the new growing meristem through symplastic connections, as a prerequisite to ED bud break. Feeding sugars into detached stems resulted in lateral bud burst and the loss of apical dominance after a few days ^11,69^. In the tuber ED system, the time gap between sugar accumulation and bud burst is suggested to be a unique mechanism. Several phytohormones are known to regulate ED in potato tubers ^17^, with ABA considered responsible for dormancy maintenance ^46^. In hybrid aspen trees, a short photoperiod signal enhances ABA level. ABA induces a pathway that enhances callose synthase blockage of PD and maintenance of bud dormancy ^47^. In maize (*Zea mays*), low temperature changed the ultrastructure of leaf PD and caused callose deposition, thereby inhibiting growth ^70^. In potato tubers, cold exposure enhances ABA ^71^. Thus, the unique time gap in ED between sugar elevation and bud burst might be due to ABA regulation of PD closure induced by cold stress. The soluble sugars induced by cold stress cannot reach the meristematic cells and catalyze growth. After the first 6 weeks, most of the soluble sugars have already been consumed, and it is therefore unlikely that sugar units induce dormancy release only via amplified energy availability. These results suggest that sugar units accumulated during ED are translated into a signal through energy sensors which induces a chain of reactions that will be resolved later in ED release.

## MATERIAL AND METHODS

### Plant material

Freshly harvested tubers of potato (*S. tuberosum* ‘Desiree’, ‘Sifra’, ‘Tyson’, and ‘Brooke’) were obtained from commercial growing fields in the northern Negev, Israel. Tubers were stored for 2 weeks at 14°C for curing. *StVInv*-knockout potato tubers were developed as previously described ^69^.

The percentage of dormancy release was measured once a week in 3 replicates of 20 tubers for each treatment. A tuber was considered non-dormant when at least one bud was 2 mm long. For sugar analysis and RNA extraction, five tubers from each treatment were sampled every 2 weeks.

### Logistic function adjustment

Dormancy-release data collected weekly during storage were used to calculate a predicted value of the logistic function with unknown *x*_50_ and *k*. Then, Excel solver was used to calculate *x*_50_ and *k*, which gave the minimum error sum of squares (SSE) for each treatment.

### Sugar extraction and quantification

Sugars were extracted and quantified as previously described, with minor modifications ^11^. Briefly, bud-base parenchyma tissue was sampled using a cork borer (Ø 1 cm, 3 cm penetration), weighed, and immediately frozen in liquid N_2_ and transferred to −80°C until use. Tissues were incubated three times in 80% ethanol at 80°C for 45 min each time. The solution was then dried using a speed vacuum (Centrivap concentrator, Labconco) and passed through a 0.2-mm membrane filter (Millex-GV filter unit, Merck Millipore). The filtrate was used for sucrose, glucose, and fructose analyses by ultrafast liquid chromatography (UFLC), using an LC-10A UFLC series system (Shimadzu) equipped with a SIL-HT automatic sample injector, pump system, refractive index detector (SPD-20A), differential refractometer detector (Waters 410) and analytical ion-exchange column (6.5 x 300 mm, Sugar-Pak I, Waters). The mobile phase (ultrapurified deionized water, Bio-Lab) was eluted through the system for 20 min at a flow rate of 0.5 mL/min, and the column temperature was set to 80°C. The chromatographic peak corresponding to each sugar was identified by comparing the retention time with that of a standard. A calibration curve was prepared using standards to determine the correlation between peak area and concentration.

### Calculation of sugar units and chilling units

To calculate sugar units, total soluble sugars were calculated as the sum (integral) of sucrose, glucose and fructose content during a selected period of time (see Eq. 2). For the chilling unit calculation, each degree below 14°C on each day of cold exposure was considered 1 chilling unit.

### Callose staining

Callose was stained as previously described, with minor modifications ^72^. Briefly, the tuber apical bud was manually detached, then sliced and incubated for 1 h in a mixture of 0.1% (w/v) Aniline blue in double-distilled water and 1 M glycine, pH 9.5, at a volumetric ratio of 2:3, respectively, and premixed at least 1 day before use. Images were acquired with a Leica SP8 laser scanning microscope equipped with a solid-state laser with 405 nm light, HC PL APO CS2 63X/1.2 water-immersion objective (Leica) and Leica Application Suite X software (LASX). Aniline blue emission signal was detected with a PMT detector in the range of 415–490 nm.

### Transmission electron microscopy

Potato tuber apical buds were detached and fixed in 3% (w/v) glutaraldehyde in 0.1 M cacodylate buffer (pH 7.4) for 3 h at room temperature, and transferred to 40°C for fixation overnight. The tissues were then rinsed four times, 10 min each, in cacodylate buffer and postfixed and stained with 1% (w/v) osmium tetroxide and 1.5% (w/v) potassium ferricyanide in 0.1 M cacodylate buffer for 1.5 h. Tissues were then washed four times in cacodylate buffer followed by dehydration in increasing concentrations of ethanol (30%, 50%, 70%, 80%, 90%, 95%) for 10 min at each concentration followed by 100% anhydrous ethanol three times, 20 min each, and propylene oxide two times, 10 min each.

Following dehydration, the tissues were infiltrated with increasing concentrations of Agar 100 resin in propylene oxide, consisting of 25%, 50%, 75% and 100% (w/v) resin, 16 h each step. The tissues were then embedded in fresh resin and polymerized in an oven at 600°C for 48 h.

Blocks with embedded tissues were sectioned with a diamond knife on a Leica Reichert Ultracut S microtome and ultrathin sections (80 nm) were collected onto 200-mesh, carbon-Formvar-coated copper grids. The sections were sequentially stained with uranyl acetate and lead citrate for 10 min each and viewed with a Tecnai 12 100 kV transmission electron microscope (Philips) equipped with a MegaView II CCD camera and Analysis^®^ version 3.0 software (Soft Imaging System GmbH).

## ACKNOWLEDGMENTS

This research was funded by a grant from the Chief Scientist of the Ministry of Agriculture and Rural Development of Israel (no. 20-06-0077). Raz Danieli’s research was supported by a fellowship from the Yair Guron scholarship fund.

## AUTHOR CONTRIBUTIONS

R.D., S.A., P.T-B., D.G., and D.E. designed the experiments. R.D., B.B.S., S.A., E.B. and Y.F performed the experiment and analyzed the data. R.D., S.A., D.G. and D.E. wrote the manuscript.

## Supplementary Figures

**Figure S1.**
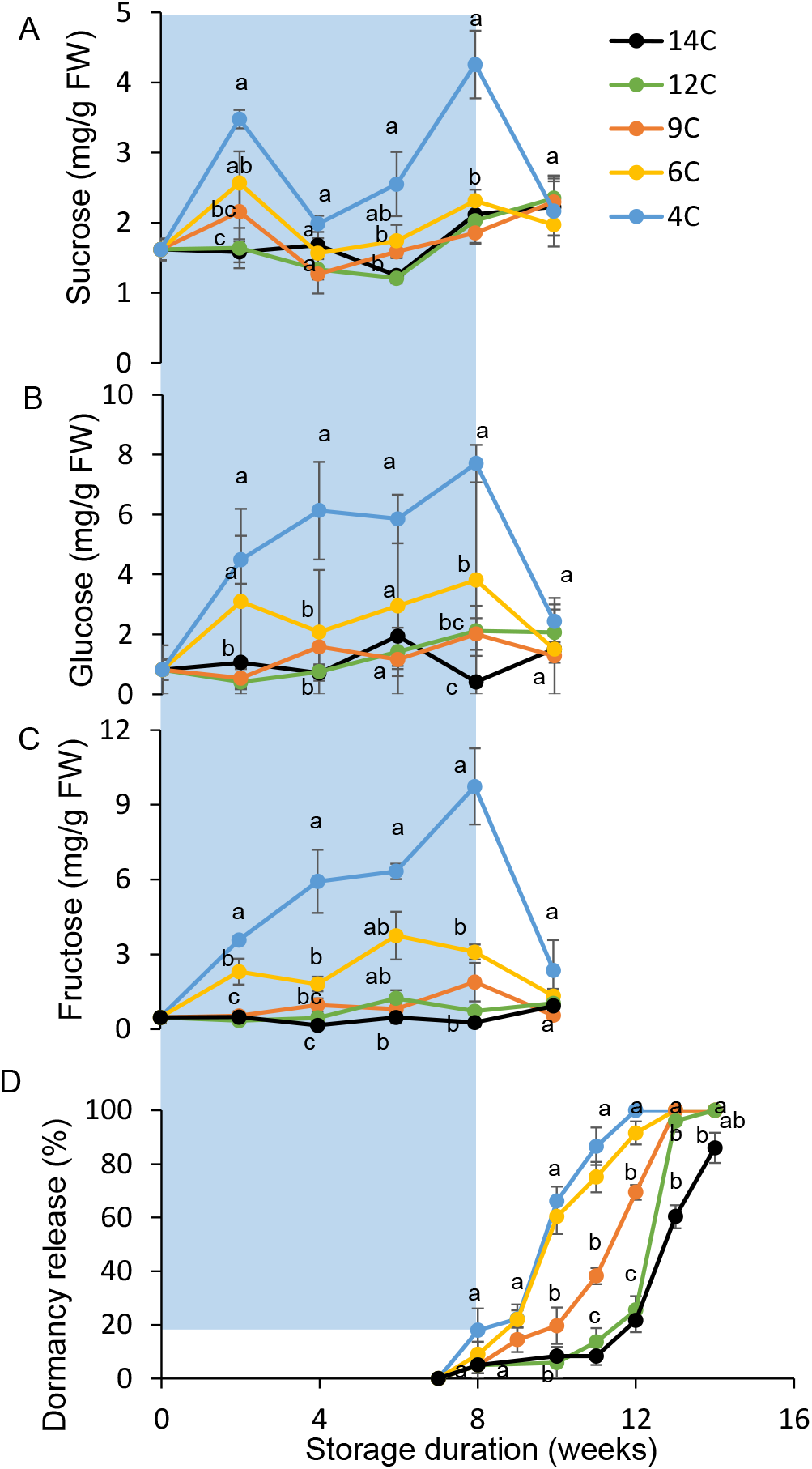
Exposure to cold storage temperature induces early endodormancy release. Desiree tubers were stored for 8 weeks (light blue background) at five different temperatures (4, 6, 9, 12 and 14°C) then transferred to 14°C storage. Every 2 weeks, five tubers from each treatment were sampled for (A) sucrose, (B) glucose and (C) fructose quantification. (D) In each treatment, three repeats of 20 tubers were measured every week for dormancy release. A tuber was considered dormant until at least one bud reached 2 mm in length. Error bars represent SE. Different letters indicate significant differences between treatments at each time point (one-way ANOVA, *p* < 0.05).

**Figure S2.**
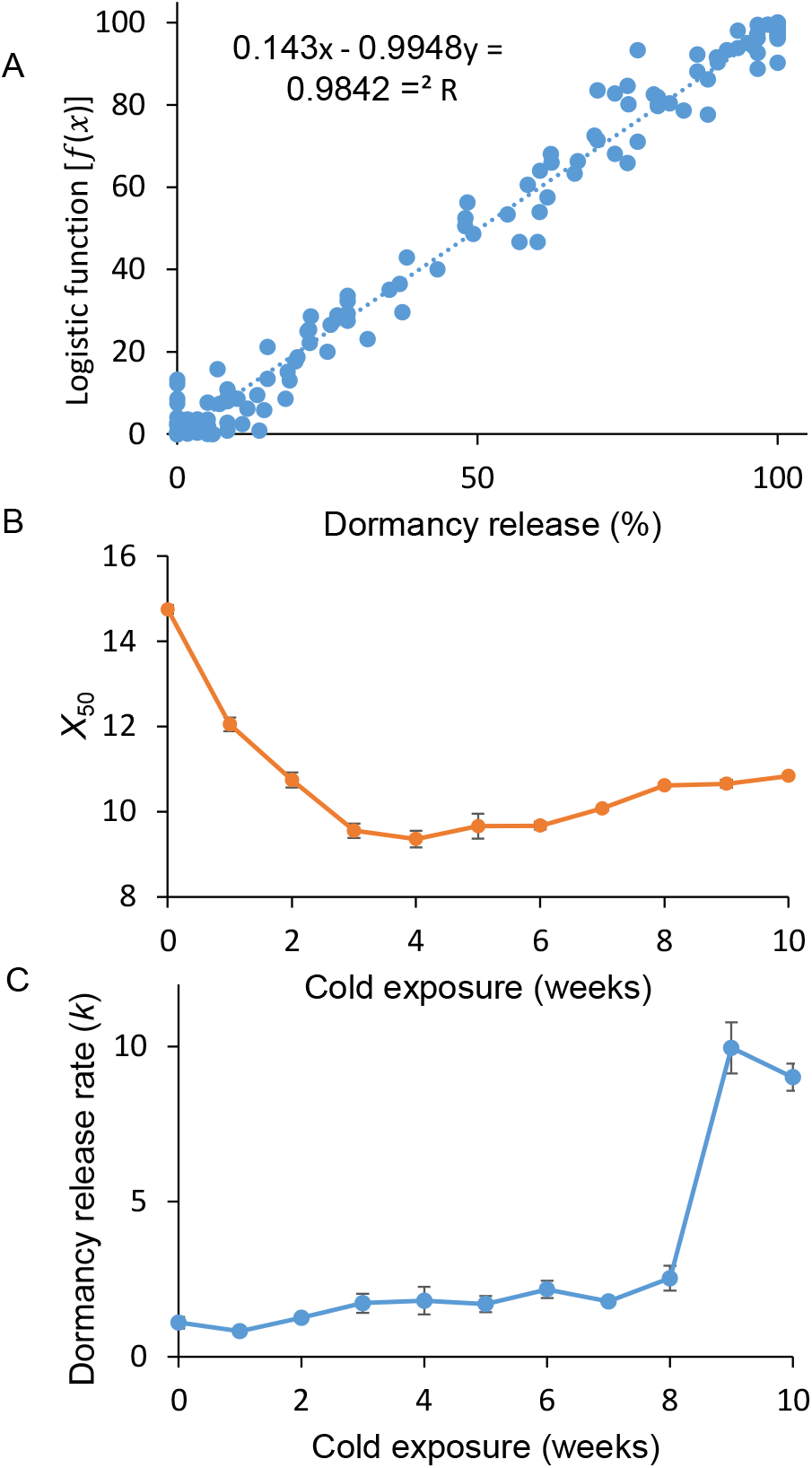
Logistic function pattern of potato tuber dormancy release. (A) Linear correlation between dormancy release phenotype and the calculated value of the logistic function. (B,C) *x*_50_ and *k* values fitted to endodormancy duration data for various durations of cold exposure. Error bars represent SE.

**Figure S3.**
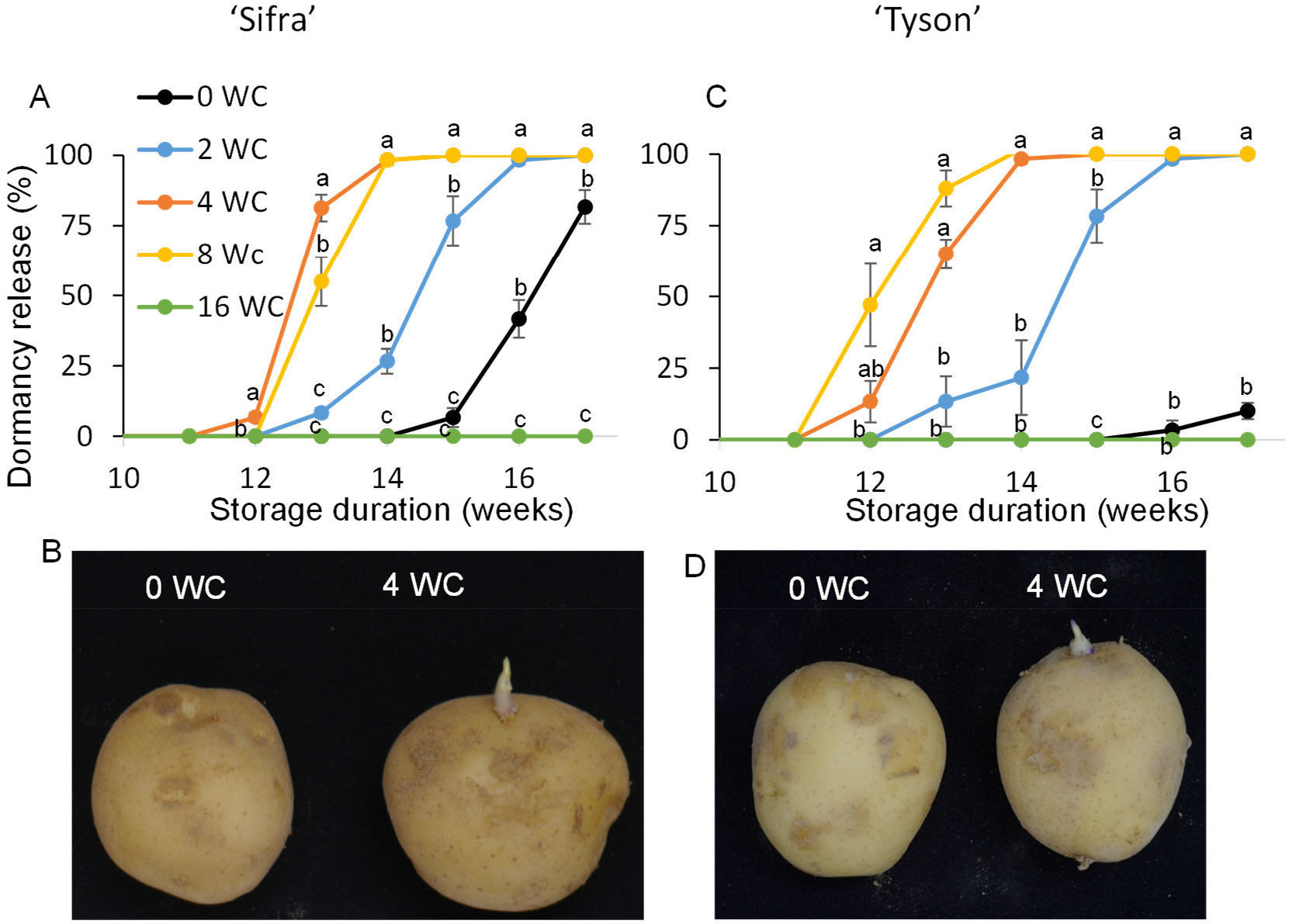
The need to absorb a sufficient amount of chilling units for ED release. ‘Sifra’ (A,B) and ‘Tyson’ (C,D) tubers were exposed to five different durations of cold (4°C) treatment (0, 2, 4, 8 and 16 weeks; WC) and then transferred to 14°C storage. In each treatment, three repeats of 20 tubers were measured every week for dormancy release (A,C). A tuber was considered dormant until at least one bud reached 2 mm in length. Error bars represent SE. (B,D) Representative images after 15 weeks of cold stress and storage.

